# Knockout of the longevity gene Klotho perturbs aging- and Alzheimer’s disease-linked brain microRNAs and tRNA fragments

**DOI:** 10.1101/2023.09.10.557032

**Authors:** Serafima Dubnov, Nadav Yayon, Or Yakov, David A. Bennett, Sudha Seshadri, Elliott Mufson, Yonat Tzur, Estelle R. Bennet, David Greenberg, Makoto Kuro-o, Iddo Paldor, Carmela R. Abraham, Hermona Soreq

**Affiliations:** The Edmond & Lily Safra Center for Brain Sciences, The Hebrew University of Jerusalem, 9190401 Jerusalem, Israel; The Alexander Silberman InsJtute of Life Sciences, The Hebrew University of Jerusalem, 9190401 Jerusalem, Israel; Rush Alzheimer’s Disease Center, Rush University Medical Center, Chicago, Illinois, USA; UT Health Medical Arts & Research Center, San Antonio, Texas, USA; Dept. Translational Neuroscience, Barrow Neurological Institute, St. Joseph’s Medical Center, Phoenix, Arizona, USA; Division of Anti-aging Medicine, Center for Molecular Medicine, Jichi Medical University, Shimotsuke, Tochigi, Japan; Dept of Neurosurgery, the Shaare Zedek Medical Center, Jerusalem; Departments of Biochemistry and Pharmacology & Experimental Therapeutics, Boston University School of Medicine, Boston, MA, USA; Klogenix LLC., Boston, MA, USA

**Keywords:** Klotho, Aging, Alzheimer’s Disease, fresh human brain, microglial and neuronal microRNAs and tRFs

## Abstract

Overexpression of the longevity gene Klotho prolongs, while its knockout shortens lifespan and impairs cognition via altered fibroblast growth factor signaling that perturbs myelination and synapse formation; however, comprehensive analysis of Klotho’s knockout consequences on mammalian brain transcriptomics is lacking. Here, we report the altered levels under Klotho knockout of 1059 long RNAs, 27 microRNAs (miRs) and 6 tRNA fragments (tRFs), reflecting effects upon aging and cognition. Perturbed transcripts included key neuronal and glial pathway regulators that are notably changed in murine models of aging and Alzheimer’s Disease (AD) and in corresponding human post-mortem brain tissue. To seek cell type distributions of the affected short RNAs, we isolated and FACS-sorted neurons and microglia from live human brain tissue, yielding detailed cell type-specific short RNA-seq datasets. Together, our findings revealed multiple Klotho deficiency-perturbed aging- and neurodegeneration-related long and short RNA transcripts in both neurons and glia from murine and human brain.

## Introduction

Age is a key risk factor for many neurodegenerative diseases^1^, and identifying transcripts distinguishing between healthy aging and age-associated cognitive decline remains a major research goal. The Klotho protein exhibits remarkable properties of lifespan regulation^2^ which attracted significant interest in aging research for over two decades. Klotho-deficient mice show premature aging symptoms, including cognitive decline^3^, whereas Klotho excess extends lifespan and improves cognitive functions^4,5^. Injected Klotho enhances cognition in aged rhesus macaques^6^, while Klotho overexpression improves amyloid-β clearance and cognition in one AD mouse model^7^, and LTP and behavior in another AD model^5^. Moreover, human AD patients show lower Klotho levels in the cerebrospinal fluid (CSF) than age-matched controls^8^, contrasting a human variant that elevates the levels of the functional Klotho protein and which associates with both longevity^9^ and lower AD risk^10^. Additionally, Klotho overexpression improved remyelination, avoided motor neurons death and lowered neuroinflammation in a multiple sclerosis mouse model ^11,12^.

While Klotho’s involvement in multiple brain processes including myelination and synaptic plasticity^5,13^ is well recognized, the impact of Klotho deficiency on long and short RNA transcriptomes is incompletely understood. To address this issue, we compared short and long RNA-sequencing profiles of brain tissues from Klotho knockout (KO) murine models to those of wild-type (WT) mice (n=5 in each group), which identified changes in numerous long and short RNA fragments. We sought correlations of the observed transcriptomic changes with reported aging and AD alterations and pursued the cell type-specificity of the observed differences. Among short RNAs, we focused on miRs and tRFs, due to their suggested involvement in brain aging and neurodegeneration^14–16^.

### Klotho knock-out induced long RNA transcriptome perturbations

Klotho deficiency perturbed 10% (1,059 out of 10,334) of all of the expressed long RNA transcripts in Klotho KO murine brains compared to WT (padj < 0.05; Fig. 1A,B, Supplementary table 1). Klotho mRNA itself was predictably downregulated in the brains of KO mice (padj<<0.001; Supplementary Fig. 1). Other differentially expressed (DE) transcripts showed smaller albeit significant changes, possibly representing the averaged expression differences between diverse cell types and brain regions in the analyzed samples. Briefly, Klotho deficiency led to impaired functioning of multiple cellular compartments, for example by modulating filamin B (Flnb), involved in cytoskeleton formation of developing neurons^17^, the Neat1 biomarker of nuclear paraspeckles^18^, Slc6a6, involved in synaptic GABA uptake^19^, and Gatm, a mitochondrial enzyme participating in creatinine synthesis^20^. Together, those differences clearly distinguished Klotho-knockout from wildtype samples in principal component analysis (PCA; Fig. 1C).

**Figure 1.**
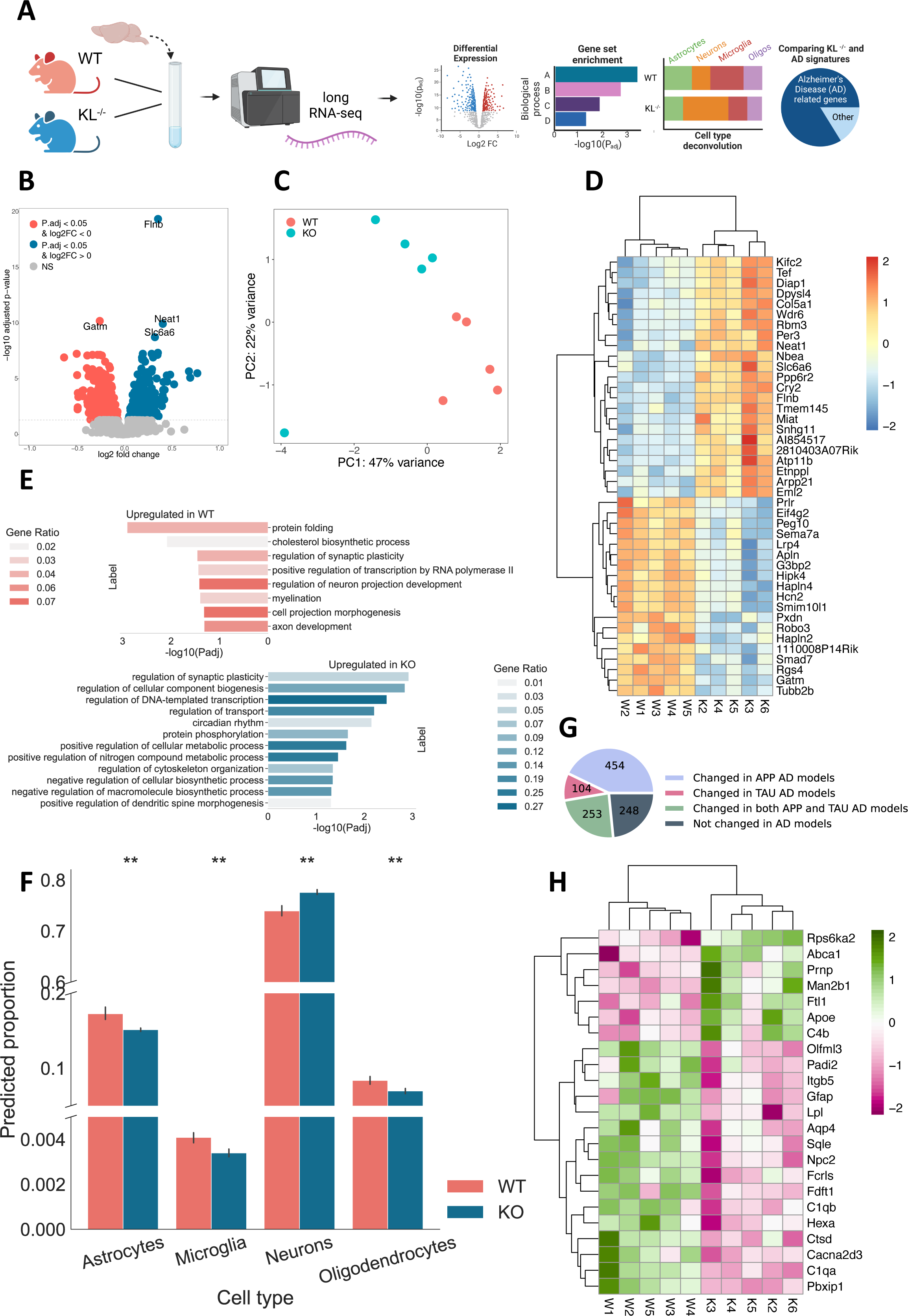
Klotho KO induces numerous long RNA transcriptome changes. **A)** Graphical representation of the research plan. **B)** Volcano plot of mRNAs DE in Klotho-knockout murine brains compared to wildtype (blue, pink: upregulated and downregulated fragments; p_adj_<=0.05). **C)** PCA based on mRNA reads colored by genotype. **D)** Heatmap showing normalized and centered counts of the most significantly altered transcripts (padj <= 5e-6; colormap - z-score of normalized count levels, centered to the mean). **E)** Gene Ontology annotation of biological processes enriched in down- and up-regulated transcripts (pink, blue) of Klotho-knockout. F) AutoGeneS prediction of the cell type proportions in WT and KO profiles based on the scRNA-seq atlas^22^; **: pval<0.01, ns: non-significant. G) Pie chart proportions of DE genes in AD studies^24^. H) Heatmap based on our Klotho KO data, showing normalized and centered counts of (G) transcripts that changed in >= 7 different AD studies.

Using the Gene Ontology resource^21^, we identified the DE genes’ biological pathways. Klotho KO reduced protein folding and phosphorylation processes, indicating disrupted protein metabolism and exacerbated protein aggregation (Fig. 1E). Myelination and neuron projection development were also decreased in Klotho KO, while genes involved in cytoskeleton organization, transport and dendritic spine morphogenesis were elevated, reflecting general morphological changes in neuronal structure (Fig. 1D,E). Considering that the analyzed bulk RNA samples represent mixtures of multiple brain cell types, we further deconvoluted the bulk RNA profiles using a single-cell RNA-seq mouse brain atlas^22^. An AutoGeneS package^23^-based cellular heterogeneity search indicated that Klotho KO caused a signaling decline in all glial cell types (Fig. 1F).

The transcriptional perturbation caused by Klotho KO further mimicked changes seen in AD-mouse models. Meta-analysis of the AD-associated transcriptome^24^ (Methods) revealed that the DE transcripts identified in the Klotho KO profiles highly overlapped with the mouse model AD-altered transcripts (Fig. 1G). Briefly, 811 out of the 1059 DE genes in Klotho KO (∼77%) were altered in at least one of the mouse models, with 454 modulated in the APP models (43%), 104 – in MAPT models (10%), and 253 in both (24%). The overlapping genes included known AD markers which changed in more than 7 different studies^24^, including the lipid metabolism genes Abca1 and Apoe^25^; complement immune system members C4b, Ctsd, C1qb and C1qa^26^, and the astrocytic markers Gfap and Aqp4^27^ (Fig. 1H; Supplementary Table 1). Thus, the transcriptional signature of murine Klotho KO shared basic features with the transcriptomic profiles associated with diverse mouse AD models.

### Klotho knock-out perturbs brain microRNA (miR) signatures

Apart from mRNAs, Klotho KO altered the levels of 27 miRs (Fig. 2A,B; Supplementary table 2), whose signatures separated the samples based on the genotype on the PCA plane (Fig. 2C). Moreover, unsupervised clustering based on the profiles of the DE miRs reliably separated the samples to genotype-specific groups (Fig. 2D). Altered miRs include mmu-miR-212-5p, linked to longevity via regulating Sirt1 and cooperating with the AD-depleted mmu-miR-132-5p^28^ (Fig. 2D). DE validated mRNA targets of the DE miRs^29^ reflect mRNA perturbations which suggest miR regulation. Identifying DE miRs targets using miRNet^30^ revealed the experimentally validated targets whose levels were repressed by the corresponding miRs in other experimental systems^31^. Further calculation of Pearson correlation coefficients between every pair of a DE miR and its DE validated mRNA target enabled comparing the correlation coefficients between the miR - target pair to pairs of miRs with non-target mRNAs (miR - non-target). Intriguingly, miR-target profiles were more negatively correlated than miR-non-target mRNA pairs (Fig. 2E), supporting the prediction that at least part of the vast mRNA perturbations following Klotho KO might be explained by miR regulation. Using the Gene Ontology resource, we have further identified DE miR target genes involved in major neuronal pathways, such as axonogenesis, dendritic spine development and regulation of synaptic plasticity (Fig. 2F). Further supporting active miRs involvement in the Klotho-KO-activated downstream processes, we found Klotho KO -elevated levels of the microprocessor-associated protein Ddx5, which is actively involved in miR processing^32^, and is targeted by four different downregulated miRs (Supplementary table 3).

**Figure 2.**
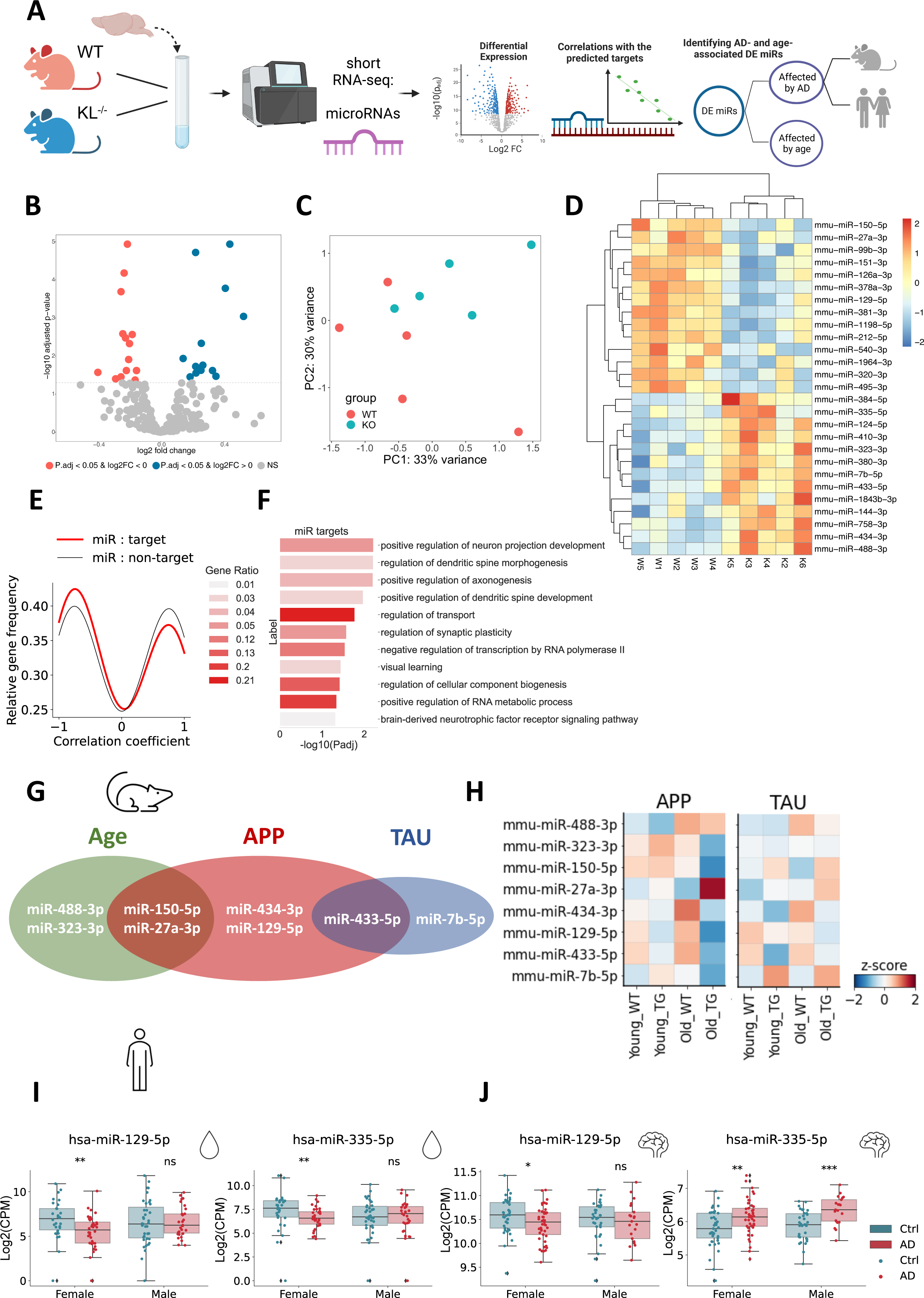
Klotho deple1on perturbs miR signatures, resembling AD and aging-associated changes. A) Graphical abstract. B) Volcano plot of miRs DE in Klotho-knockout compared to wildtype mice (blue, pink: upregulated or downregulated fragments; p_adj_<=0.05). C) PCA based on miR counts colored by genotype. D) Gaussian kernel density estimates of Pearson correlation coefficients between klotho-affected miRs and their targets (red), and klotho-miRs and DE mRNAs which are not their targets (black). X axis: Pearson correlation coefficient. Y axis: relative frequency of miR-mRNA pairs with the corresponding coefficient. E) Heatmap showing normalized and centered counts of DE miRs (colormap: z-score of the normalized counts, centered to the mean). F) Gene Ontology annotation of miR targets from enriched biological processes. G) Venn diagram of DE miRs that co-change in the murine hippocampus with age and/or in one of the AD pathology models^33^ (APP - APP^swe^/PS1^L166P^; TAU - THY-Tau22). H) Scaled heatmap of miR alterations from (G) across age and genotype conditions of both APP and TAU mouse models^33^. I) Log_2_(CPM) counts of hsa-miR-129-5p and hsa-miR-335-5p in AD patients’ CSF and cognitively healthy controls, divided by sex^36^ (**: p_adj_ <0.01). J) Log_2_(CPM) counts of hsa-miR-129-5p and hsa-miR-335-5p in the Nucleus Accumbens of AD patients (cogdx=4), compared to cognitively healthy individuals (cogdx=1), divided by sex^16^ (**: p_adj_<0.01).

To find out if Klotho DE miRs are also altered in aging or AD pathology, we analyzed short RNA-seq data from AD mouse models, including young and aged (4 and 10 months) wildtype and transgenic mice carrying either APP (APP^swe^/PS1^L166P^) or TAU (THY-Tau22) mutations^33^. For each strain, we compared miR profiles from wildtype and age-matched transgenic mice, and along age within the same genotype (Supplementary Table 4). This identified Klotho-associated miRs whose levels were affected by age (e. g. miR-488-3p; miR-323-3p), or by both age and APP pathology (miR-150-5p, miR-27a-3p), by APP pathology alone (miR-434-3p, miR-129-5p), by TAU pathology alone (miR-7b-5p), or by both APP and TAU pathologies (miR-433-5p; Fig. 2G,H), together reflecting altered regulation of cognition.

Next, we sought putative human links between DE miRs, cognition and AD neuropathology. Intriguingly, two miRs DE in Klotho KO, miR-129-5p associated with the Fragile X syndrome^34^ and the muscarinic receptors-targeting miR-335-5p^35^, were reduced in the cerebrospinal fluid (CSF) of female, but not male AD patients^36^ (Fig. 2I). Considering our recent findings of cognition-related female-specific AD declines of tRF regulators of cholinergic mRNAs^24^, this suggests links between Klotho-KO-induced transcriptomic perturbations to sex differences in dementia. Indeed, both miR-129-5p and miR-335-5p are also altered in the Nucleus Accumbens of female AD patients, compared to controls with no cognitive impairment^16^ (Fig. 2J; Supplementary Table 5), and with the same direction of change in human AD and murine Klotho KO brains. The down-regulated transcripts and processes hence emerge as co-related to the cognitive impairments shared by Klotho KO and AD.

### Klotho knockout alters the levels of short tRNA fragments (tRFs)

Apart from DE miRs, we also identified Klotho-associated expression changes in another short non-coding RNA family – tRFs^11^ (Fig. 3A). Six nuclear genome-originated tRFs were altered, three upregulated and three downregulated (Fig. 3B; Supplementary table 5). Strikingly, all three upregulated tRFs were 3’ halves originated from Asparagine-GTT tRNA, representing matching 3’-end sequences with single nucleotide differences, whereas the three downregulated ones were Leucine-TAG tRNA-originated i-tRFs (Fig.3B; Supplementary table 5). These subtle differences allowed PCA to separate the samples (Fig. 3C), and unsupervised hierarchical clustering based on the DE tRFs recognized the knockout and wildtype genotypes as two distinct clusters (Fig. 3D).

**Figure 3.**
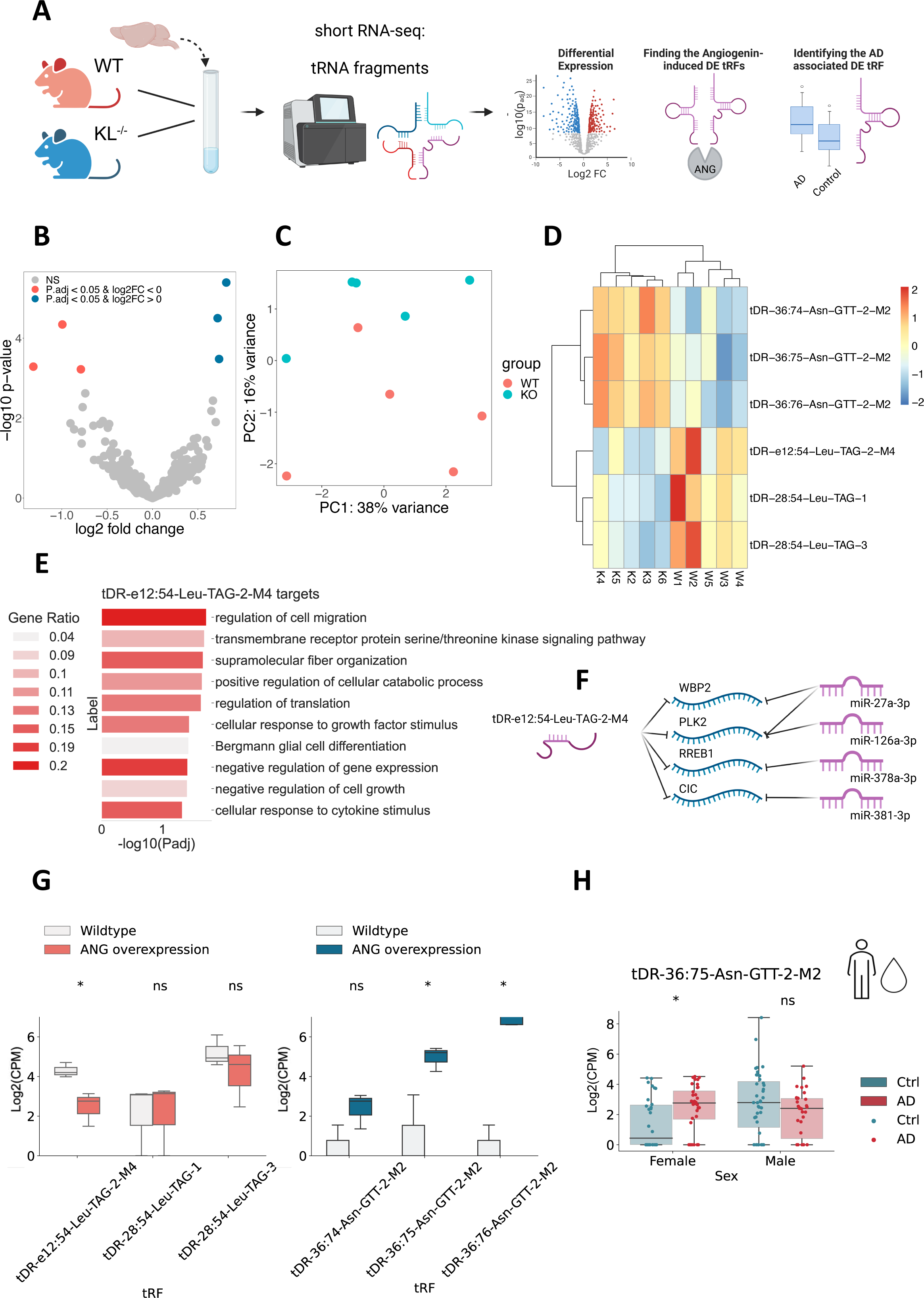
Klotho knockout alters tRF levels. A) Graphical abstract. B) Volcano plot of tRFs DE in Klotho-KO mice compared to wildtype; up- and downregulated (blue, pink); p_adj_<=0.05). C) PCA of tRF counts colored by genotype. D) Heatmap showing normalized and centered counts of altered tRFs (colormap: z-score of the normalized counts, centered to the mean). E) Gene Ontology annotation of biological processes enriched among tDR-36:75-Asn-GTT-2-M2 targets, negatively correlated with this tRF in Klotho KO data. F) Schematically illustrated common targets of tDR-36:75-Asn-GTT-2-M2 and miRs. G) Log_2_(CPM) counts of altered tRFs in HEK293 cells under Angiogenin overexpression compared to wildtype; le_, right – tRFs declined or elevated under Klotho KO; ^38^ (*: p_adj_ <0.1). H) Log (CPM) counts of tDR-36:75-Asn-GTT-2-M2 in the CSF of AD patients and cognitively healthy controls, divided by sex^36^ (*: p_adj_ <0.05).

Out of all of the DE tRFs, only tDR-e12:54-Leu-TAG-2-M4, downregulated in Klotho KO, was short enough to predict miR-like activity^15^. The tRFTar tool^37^ identified 81 predicted mRNA targets that were negatively correlated with tDR-e12:54-Leu-TAG-2-M4 (Supplementary table 6). Notably, these tRF targets were enriched in distinct biological processes from those enriched by the Klotho-altered miR targets, including cell migration, intracellular signaling and translation regulation pathways (Fig. 3E). Moreover, tDR-e12:54-Leu-TAG-2-M4 shared 4 targets with downregulated miRs (Fig. 3F), possibly reflecting cooperation of these short non-coding RNA families in knockout-induced transcriptional changes.

The transcriptomic profiles further revealed slightly upregulated levels of the nuclease angiogenin mRNA linked to tRF production in Klotho KO (Supplementary Fig. 2B)^38^. To determine whether those DE tRFs were stress-induced products of angiogenin^38,39^, we sought parallel tRF changes in relevant model systems. Noticeably, Klotho KO shared all of its upregulated tRFs with angiogenin overexpression in HEK293 cells, and one of the downregulated tRFs, tDR-e12:54-Leu-TAG-2-M4, decreases in angiogenin overexpression (Fig. 3G) and increases in U2OS cells under angiogenin knockout^38^ (Supplementary Fig. 2A). Thus, the tRF signature following Klotho knockout might represent a stress-induced and angiogenin-dependent response.

Comparing the identified tRF changes under Klotho knockout to those of AD brains identified tDR-36:75-Asn-GTT-2-M2, as both upregulated under Klotho knockout and in the CSF of female AD patients, compared to cognitively unimpaired controls^36^ (Fig. 3H). Mimicking the sex-specific miR changes and decline of tRFs in the nucleus accumbens from AD female brains^16^, tDR-36:75-Asn-GTT-2-M2 levels remained unchanged in male patient brains. In summary, both Klotho knockout-induced brain miRs and tRFs changes correspond to their parallel changes in AD pathology.

### Klotho-miR profiles reveal neuronal and microglial signatures

Deconvoluting the bulk mRNA signal to cell type-specific signatures showed that Klotho KO led to reduced numbers of glial transcripts (Fig. 1F). However, deconvoluting the affected miRs, which are highly cell type-specific^40^ was challenging due to the lack of reference datasets with cell type resolution. Therefore, we developed novel neuron- and microglia-specific short RNA-seq datasets from live human brain samples resected during surgeries for the removal of non-infiltrative brain tumors. To assess the non-tumor origin of the cortical tissues, we excluded patients with infiltrative brain pathologies, such as meningitis, glioma, or diffuse cerebral inflammation, collected the tissues as far from the tumor as possible and validated that the acquired miR profiles differ from specimens of meningioma^41^, but rather resemble postmortem bulk miRNA profiles from various human brain regions^42^ (Supplementary Figure 3).

Following our inhouse NuNeX protocol^43^, the collected 16 live human brain samples (6 females, 10 males; 42-75 years old) were exposed to a brief Formalin fixation and separated into single nuclei suspensions which retained some associated cytoplasm. Staining with neuron-specific (NeuN) and microglia (Iba1) fluorescent markers followed by FACS sorting yielded 32 microglia- and neuron-specific populations and their short RNA-seq profiles (Fig. 4A; Supplementary tables 6, 7). Since Iba1 is also expressed by macrophages that infiltrate the brain^44^, we validated the microglial identity of the sorted Iba1^+^ populations using qPCR measurement of specific microglial RNA markers, such as P2y12, Sall1, Tmem119 (Supplementary Figure 4).

**Figure 4.**
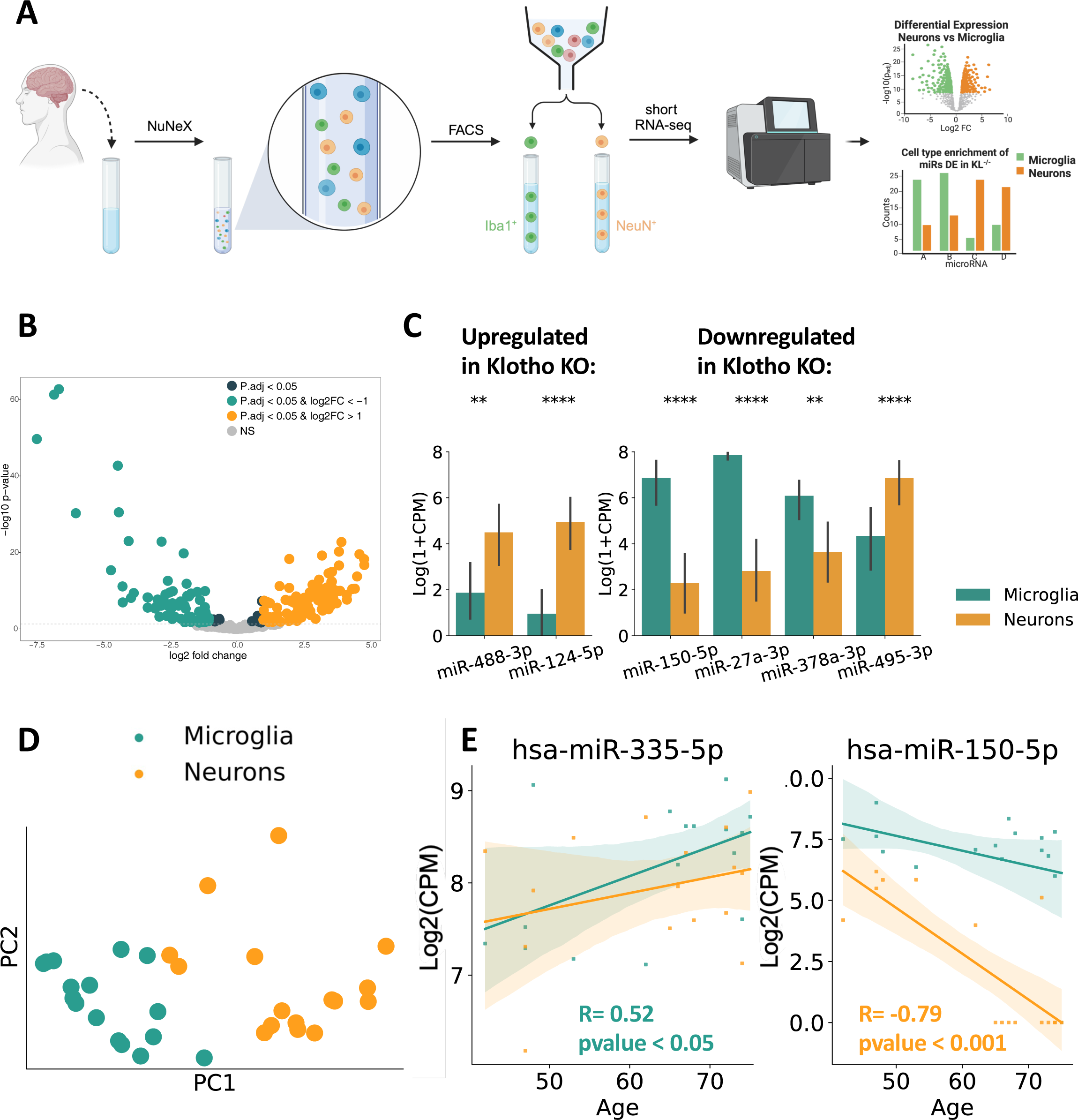
Klotho KO affects miR levels in both neurons and microglia. A) A schematic pipeline used to produce bulk cell type-specific short transcriptome datasets of live brain cells. Using the NuNeX protocol^43^, samples extracted during neurosurgeries were broken down to single nuclei surrounded by thin cytoplasm layers. Nuclei stained with microglia-(Iba1) and neuron-(NeuN) specific fluorescent markers were FACS-sorted and short RNA-sequenced. B) Volcano plot showing altered level miRs enriched in neurons or microglia (green, orange). C) 6 miRS DE in Klotho KO were also enriched in one of the cell types. LeX, right: miRs upregulated or downregulated in Klotho KO, **: p_adj_<=0.01; ****: p_adj_<=0.0001. D) PCA of the cell type-specific small transcripts (7 miRs DE in Klotho KO, colored by cell type). E) Linear correlation of operated donors age. Log_2_(CPM) count levels of hsa-miR-335-5p (correlated in microglia) and hsa-miR-150-5p (correlated in neurons), colored by cell type.

Intriguingly, our novel short RNA-seq datasets revealed highly distinct miR profiles between live human brain neurons and microglia (Fig. 4B). Using the DESEQ2 package^45^, we found that 57% of the expressed miRs (173 of 303) were enriched in one of these cell types (Supplementary table 8). Among them, 69 and 91 were upregulated in microglia or neurons with a log-fold change higher than 1, yielding highly valuable resources for deconvolution of bulk brain-derived short RNA profiles.

Of the 27 DE miRs in murine Klotho KO, 7 were found in our newly produced cell type-specific human dataset, with 6 of those selectively enriched in one of those cell types (Fig. 4C; Supplementary table 8). Parallel to the decline in glial mRNA transcripts under Klotho KO, 3 out of 4 miRs declined in Klotho KO were enriched in human brain microglia (Fig. 4C). In contrast, miRs upregulated in Klotho KO were enriched in human neurons from live human brain samples, mimicking the elevated neuron-specific mRNA transcripts in our mRNA-based deconvolution. Further, the cell type-dependent expression of the 7 DE miRs in Klotho KO sufficed to separate microglial and neuronal samples, showing a strong cell type-specific signal in the Klotho KO miR profiles (Fig. 4D) with shared cell type specificity of the DE miRs seen between the sexes (Supplementary Fig. 5).

Notably, two DE miRs in Klotho KO showed cell type-dependent age correlation (Fig. 4E). In particular, the cholinergic miR-335-5p, which targets the muscarinic receptors and might limit the cholinergic blockade of inflammation^35,46^, was upregulated with age in human brain microglia, murine Klotho KO and AD patients’ CSF and Nucleus Accumbens (Fig. 2I,J). In contrast, miR-150-5p, which targets the autism-associated Foxp1^47^ transcript, declined with age in live human neurons, Klotho KO and by both age and APP pathology in AD mice models (Fig. 2G,H). Thus, our findings further strengthened the notion of miR-335-5p and miR-150-5p contributions to aging-related processes.

Notably, aligning our neuron- and microglia-specific short RNA datasets to a tRF database (Methods) produced a novel RESOURCE of cell type-specific tRFs from live human brains. Unlike miRs, there were many more tRFs elevated in neurons than in microglia (Supplementary Fig. 6A). While none of the tRFs DE in Klotho KO were expressed in human neurons or microglia, other fragments originating from the same tRNAs exhibited a trend of elevation in neurons, compared to microglia (Supplementary Fig. 6B), possibly indicating that tRFs signaling is generally more potent in neurons than in microglia. The neuron-specific tRF dataset is openly available (Supplementary table 9).

This study has potential limitations. First, our discovery of overlapped molecular signatures between Klotho KO, human brain aging and AD is correlative and needs to be validated experimentally. Furthermore, our novel cell type-specific short RNA-seq dataset from live human brain tissues is prone to be biased by the pathology effect. We showed that the extracted profiles are closer to those of healthy postmortem brains than to meningioma samples (Supplementary Figure 3), similar to multiple previous studies on live human brain tissue, which suffered from the same limitation^48–50^. However, we cannot fully rule out the possibility that the extracted neuronal and microglial profiles are affected by the tumor.

### Conclusions

Klotho is an anti-aging gene with a promising therapeutic potential^51^. To seek mechanisms by which Klotho exerts its CNS effects, we compared the long and short RNA profiles of murine Klotho-deficient brains with those of healthy controls. This revealed a major Klotho knockout - induced transcriptional dysregulation affecting multiple cellular compartments and brain cell types. Importantly, many Klotho KO-altered long and short RNA fragments were also changed in brain aging and diverse AD-related pathologies, possibly reflecting Klotho’s capacity to ameliorate aging-related cognitive decline. Furthermore, deconvolution analysis using a single-cell RNA-seq atlas of the murine brain^22,23^ showed that Klotho knockout causes a slight decline in glial mRNA transcripts. By generating a novel cell type-resolved short RNA dataset from live human brain tissues, we further revealed that Klotho KO led to reduced levels of microglia-enriched miRs while elevating the levels of neuronal miRs. Thus, the general transcriptional effect of Klotho knockout in the murine brain provides a key RESOURCE for aging researchers with a focus on Klotho-associated brain processes. Moreover, the data presented here indicates that physiologically normal Klotho levels are necessary for white and gray matter development and maintenance during life. Since Klotho levels decrease with age, augmenting Klotho protein expression either by enhancing its endogenous expression or by delivering exogenous genetic material, would be beneficial against neurodegenerative and other age-related diseases.

## Methods

### Klotho KO mice

#### RNA extraction

WT and Klotho knock-out (KL^-/-^) female mice, n=5 in each group, were sacrificed at 6 weeks of age by cervical dislocation and the whole brain removed and snap-frozen in liquid nitrogen before storage at −80°C. Each brain was dissected on dry ice into roughly ten pieces which were placed in QIAzol (Qiagen, 217004) and immediately homogenized using a pellet pestle. All homogenates of each brain were then pooled and homogenized again using a Polytron PT 3000 (Kinematica) and stored at −80°C. RNA was extracted from a 700 µL aliquot of each homogenate using the miRNeasy Mini Kit (Qiagen, 217004) according to the manufacturer’s protocol. RNA concentration was determined (NanoDrop 2000, Thermo Scientific) and RIN was measured (Bioanalyzer 6000, Agilent), with all samples ranging from 8.3 to 8.9.

#### RNA sequencing and alignment

All samples underwent small RNA-sequencing and poly(A)-selected long RNA-sequencing. For small RNA, libraries were constructed from 1000 ng total RNA (NEBNext Multiplex Small RNA library prep set for Illumina, NEB-E7560S, New England Biolabs) and the small RNA fraction sequenced (NextSeq 500/550 High Output Kit v2.5 75 Cycles, 20024906, Illumina). For poly(A)-selected RNA, libraries were constructed from 1000 ng total RNA (KAPA Stranded mRNA-Seq Kit,KR0960–v5.17, Kapa Biosystems) and the long RNA sequenced (kit as above). All new sequencing was done on the NextSeq 500 System (Illumina) at the Center for Genomic Technologies Facility, the Hebrew University of Jerusalem. Long RNA was aligned to the human reference (ENSEMBL GRCh38 release 79) using STAR^52^. Small RNA was aligned to miRBase using miRexpress^53^ and to the tRNA transcriptome using the MINTmap^54^. The nomenclature of tRFs was based on the recently published standardized ontology^55^.

### Live human brain samples

#### Tissue sampling

The study, as well as acquisition of cortical brain samples, was approved by the institutional review board (IRB protocol RMB-0713-19) and by the national ethical review board. All patients signed informed consent forms. The biopsies were extracted in operations for resection of non-neural pathologies, including meningiomas, metastases and vascular malformations. We excluded patients who were operated for neural pathologies (e. g. gliomas). We noted patient age, sex, the pathology for which the patient was operated, and whether or not edema was noted in the biopsied brain specimen. We also noted and evaluated other parameters relevant to neural tissue, microglia or both, including previous brain irradiation, past medical history, current medications, monocyte counts and fractions and others. In the operating room, samples were collected from the cortex overlying the tumor. Sampled brain tissue was fixed in formaldehyde 4% (Sigma-Aldrich HT501128) for 30 minutes. The samples were washed with PBS and then stored in RNAlater solution (Merck; R0901-100M) until homogenization.

#### NuNeX

Samples were homogenized with a Dounce tissue grinder (Sigma-Aldrich, D9063). The homogenate was passed through a 40 μm cell strainer (BD Falcon) and pelleted by centrifugation at 900 *g* for 5 min. Cells were resuspended and stored in −80°C until FACS sorting.

#### Immunostaining and FACS

Before cell sorting, Fc block (Invitrogen, Waltham, MA, USA,14-9161-73) was added according to manufacturer’s instructions at 20 μL/tube and incubated at 4°C for 20 min. For staining, anti-NeuN (1:500, Alexa Fluor^®^488 conjugated Sigma #MAB377X) and anti-Iba1 (1:500; Abcam #ab178846) antibodies were added for 30 min incubation. Cells were then pelleted by centrifugation at 900 *g* for 5 min, resuspended in staining buffer, and stained with the secondary antibody for anti-Iba1 (1:500; Alexa Fluor 647 conjugated Jackson #711-605-152), followed by 30 min incubation. Re-pelleted cells were resuspended in staining buffer with DAPI (1:1000; Santa Cruz, sc-3598). All steps were performed on ice. Iba1- and NeuN-positive cells were sorted through a 85 μm nozzle with an approximate flow rate of 8,000 events/s. Sorted Iba1- and NeuN-positive cells were collected into tubes containing 500 μl staining buffer (> 1,000 cells), then centrifuged at 900 *g* for 5 min and resuspended in 100 μl PKD buffer. RNA was extracted using a specialized kit for RNEasy FFPE Kit (Qiagen, cat. No 73504). Samples were collected with BD FACSAria III (BD Biosciences) and analyzed with FCS Express 7 SoXware (*De Novo* SoXware).

#### RNA sequencing and alignment

All samples underwent sequencing of small RNA. Libraries were constructed from 500 pg total RNA using D-Plex Small RNA-seq Kit for Illumina and Single Indexes for Illumina - Set B (Diagenode, C05030001). RIN aXer cell sorting was determined to be between 1.2 and 7.2 (Bioanalyzer 6000, Agilent) and the small RNA fraction was sequenced using NextSeq 2000 P3 Reagents (20046810, Illumina) on the NextSeq 2000 System (Illumina) at the Center for Genomic Technologies Facility, the Hebrew University of Jerusalem. Small RNA was aligned to miRBase using miRexpress^53^ and to the tRNA transcriptome using the MINTmap^54^.

### ROSMAP subjects

ROSMAP Subjects: RNA-Sequencing (RNA-Seq) data was derived from participants in the Rush Alzheimer’s Disease Center’s ROS and MAP cohort studies^42^. Both studies were approved by an Institutional Review Board of Rush University Medical Center. All participants in ROSMAP enroll without known dementia and agree to annual clinical evaluation and brain donation at the time of death. Subjects are clinically classified for dementia, AD and Mild Cognitive Impairment (MCI) as previously reported^56,57^. Those without dementia or MCI were designated as no cognitive impairment (NCI)^58^. A summary diagnosis was made by a neurologist after death based on select clinical data blinded to all neuropathologic data. All cases received a neuropathological evaluation based upon Braak stage and CERAD which are combined to create NIA-Reagan pathologic criteria for AD^59^. Analysis shown in Fig. 2J includes 141 *postmortem* samples from the Nucleus Accumbens. The labels (AD and control) were assigned based on the summary diagnosis of AD and NCI. Persons with MCI were excluded. The summary metadata table is provided in Supplementary Table 5.

### Data Analysis

#### Differential Expression (DE) analysis

DE analysis (Figures 1B, 2B, 3B, 4B) was performed in R using the DESEQ2 package^45^. Prefiltered raw counts of long RNAs and miRs included fragments with median expression above 10 counts per million (CPM). The raw counts of tRFs were normalized using the DESEQ2 median of ratios method in which counts are divided by sample-specific size factors determined by median ratio of gene counts relative to geometric mean per gene^45^ and prefiltered to include fragments with median expression above 10 counts.

#### Gene set enrichment

Gene set enrichment to identify biological processes (Figures 1E, 2F, 3E) was performed with the PANTHER tool of Gene Ontology resource^60^. The list of all fragments expressed above (median CPM >=0) was used as a reference gene list.

#### Cell type deconvolution

Cell type deconvolution shown in Fig. 1F was performed in python using the AUTOGENES package^23^ (Aliee et al, 2021), using a single-cell dataset from whole murine brain as a reference^22^.

#### Elucidating AD-related long RNA

Identifying AD-associated transcripts among DE long RNA (see Fig. 1G) was performed using the meta-analysis resource^24^, combining lists of DE genes in 15 AD-APP mouse model studies and 12 AD-MAPT mouse model studies. The overlap of DE genes in Klotho KO with the DE genes in APP and MAPT models was calculated separately.

#### miR and tRF target prediction

Predicted miR mRNA targets were identified using miRNet^30^, including only experimentally validated targets. Predicted targeted tRFs were identified using tRFTar^37^.

#### Elucidating AD related miRs

AD-associated miRs (see Fig. 2G), were identified using the analysis of the short RNA-seq data derived from AD mouse models^33^. MiRs, changed with age or one of AD mutations, were identified using the Wilcoxon signed-rank test, applied to log-transformed normalized counts. Fragments with p_adj_<=0.05 aXer FDR correction were considered significant.

## Acknowledgements

The authors thank the study participants of the Religious Orders Study (ROS) and Rush Memory and Aging Project (MAP) and staff of the Rush Alzheimer’s Disease Center in Chicago and express their gratitude to Dr. Ola Karmi from the Interdepartmental Instrumentation Facility (Tzabam; Hebrew University of Jerusalem) for her guidance and assistance in FACS sorting, as well as to Ms. Adi Tujerman from the Center for Genomic Technologies (Hebrew University of Jerusalem) for her assistance with RNA-seq experiments and Dr. Taylor Schmitz from the University of Western Ontario for inspiring discussions. This work was supported by the Israel Science Foundation (ISF; 1016/18; 3213/19 to H. Soreq), the National Institute of Health (NIH; Aging grant 5P01AG014449-21 to E. Mufson and H Soreq), Keter Holdings (to H. Soreq), Klogenix LLC (Boston, MA, USA), the Gatsby Charitable Foundation (to Y.L.), the K. Stein foundation (to H.S) and a joint grant by the Shaarei Zedek Medical center (to I. Paldor and H. Soreq). The ROS-MAP projects were supported by grants from the National Institute of Health (NIH; P30AG10161, P30AG72975, R01AG15819, R01AG17917, U01AG46152, U01AG61356). Serafima Dubnov is an awardee of PhD fellowships by Azrieli and Kaete-Klausner foundations. The graphical abstracts in this article were created with BioRender.com.

## AbbreviaFon list

AD: Alzheimer’s Disease
APP: amyloid precursor protein
FACS: fluorescence-activated cell sorting
FFPE: formalin fixed paraffin-embedded
Flnb: filamin B
Gatm: glycine amidinotransferase
KO: knockout
i-tRF: internal tRF type
MAP: Rush Memory and Aging Project
MAPT: microtubule associated protein tau
miR: microRNA
mmu-miR: miR from the *Mus musculus* genome
Neat 1: nuclear enriched abundant transcript 1
NFT: Neurofibrillary tangles
PBS: phosphate buffered saline
PKD buffer: protein kinase digestion buffer
tRF: tRNA fragment
RADC: Rush Alzheimer’s Disease Center
ROS: Religious Orders Study
Slc6a6: solute carrier family 6 member 6
tRFs: transfer RNA fragments
WT: wildtype

