## Supplementary Figures for "Knockout of the longevity gene Klotho perturbs aging- and Alzheimer’s disease-linked brain microRNAs and tRNA fragments"

**Supplementary Figure 1: Validation of Klotho KO**

Counts per million (CPM) of the Klotho gene in Klotho-knockout and wildtype mice

(***: $p << 0.001$).

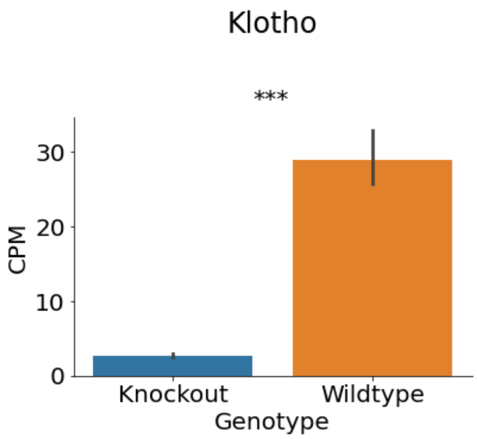


**Supplementary Figure 2: Klotho-associated tRFs might be induced by Angiogenin stress response**

A) Log(CPM) counts of U2OS cells in the Angiogenin Knockout condition and wildtype (data from Su et al, 2019; *: $p_{adj} < 0.1$). B) Log(CPM) counts of the Angiogenin gene in Klotho-knockout (KO) and wildtype mice (WT); ns – not significant.

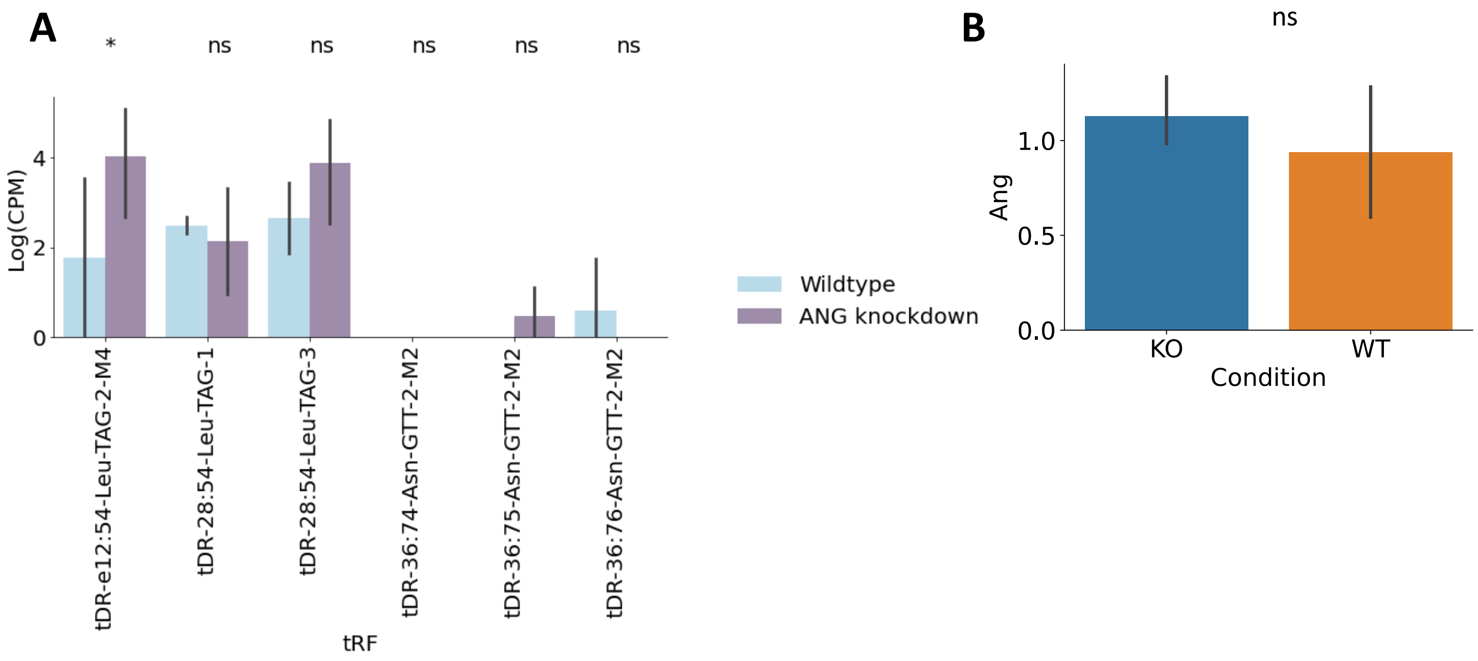

### Supplementary Figure 3: Comparing microRNA profiles of NuNeX, meningioma and postmortem brain samples

A) UMAP visualization of microRNA profiles from Meningioma[41] NuNeX and postmortem brain samples from Hypothalamus, Nucleus Accumbens (NucAcc) and Superior Temporal Gyrus (STG)[42]. NuNeX data described in this study is closer to other brain samples, in particular to cortical STG, than to meningioma. B) Correlation matrix showing pearson coefficients between the 50 averaged principal components for each cell type. C) Number of overlapping differentially expressed genes (wilcoxon test, one-vs-all comparisons, $log_2$foldchange>0, $p_{adj}$<=0.05) between the cell types.

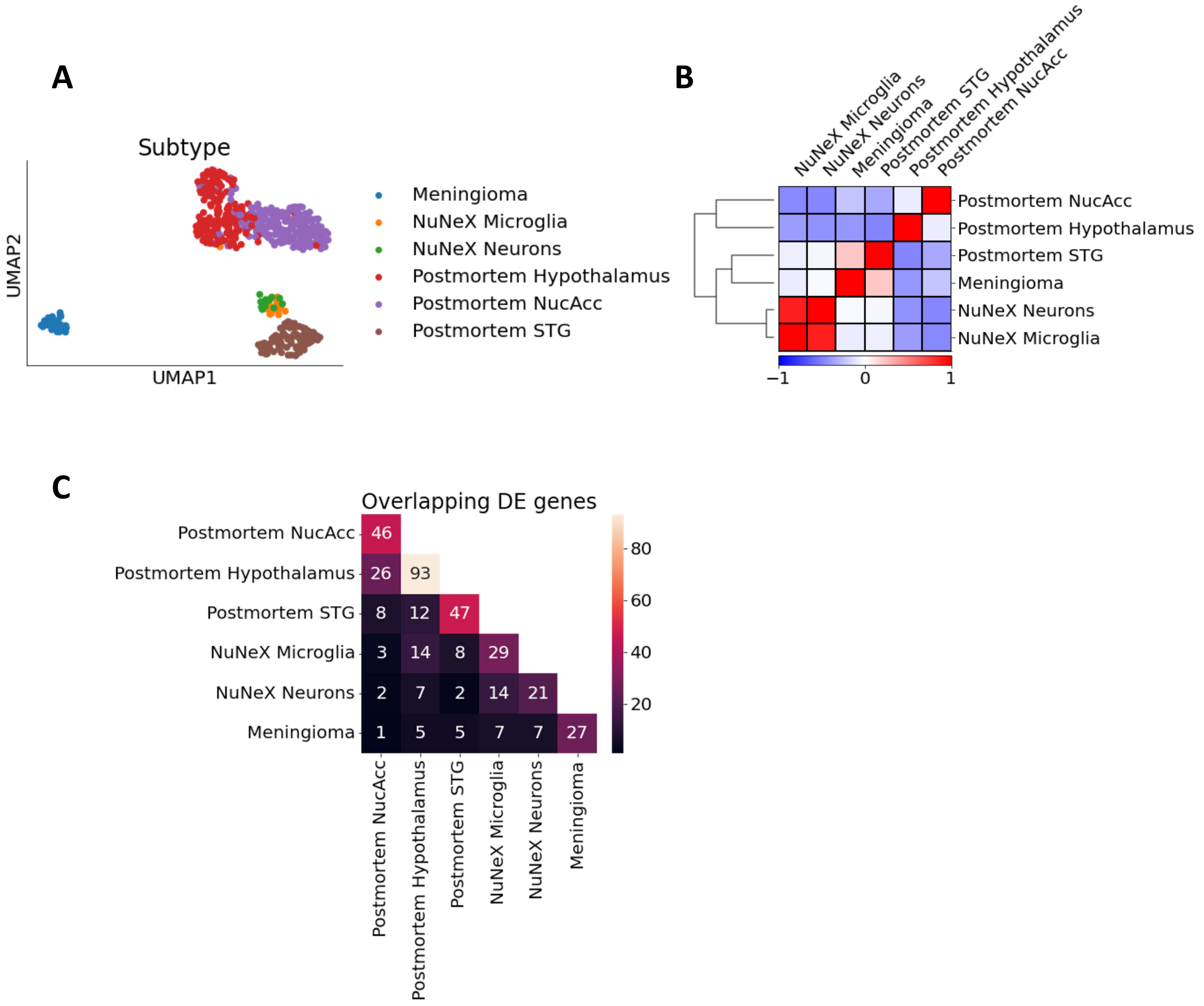

**Supplementary Figure 4: Validation of the microglia identity of the IBA1⁺ sorted population**

Change in qPCR cycles of detection of microglia (P2Y12, TMEM119, SALL1) and macrophage (SIGLEC1, CD44) markers between Iba1⁺ and NeuN⁺ sorted populations.

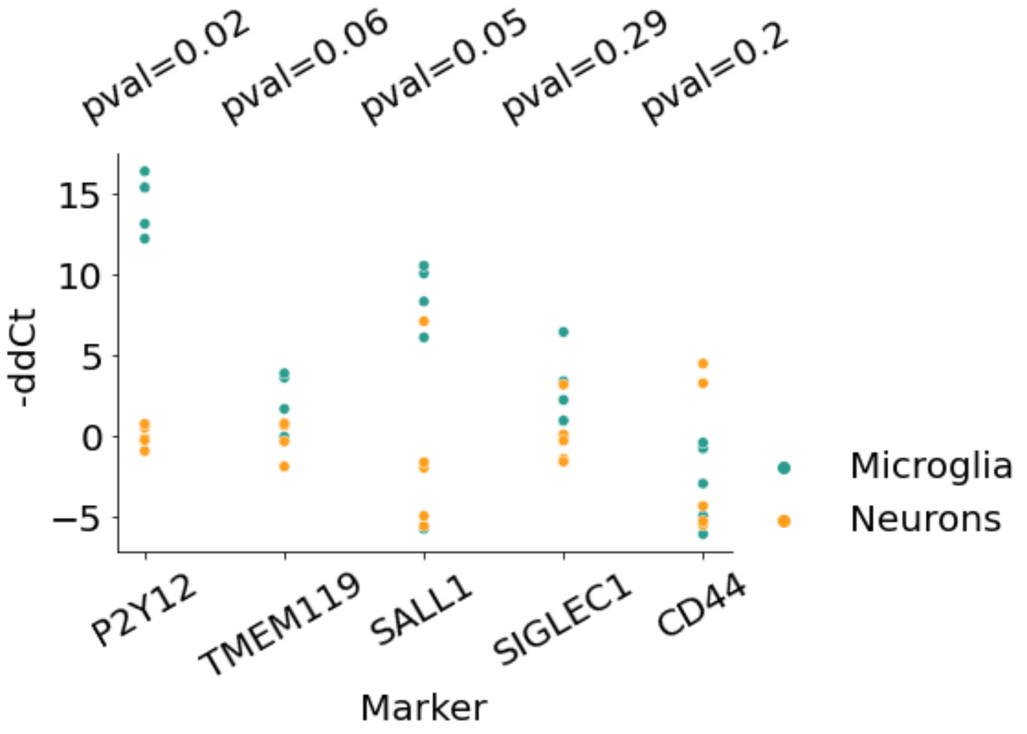


**Supplementary Figure 5: The cell type signal of miRs DE in Klotho is shared between the sexes**

A. PCA based on microRNA profiles of FACS-sorted neuronal and microglial populations, colored by sex. B) Same as (A) but only from female samples, colored by cell type. C) Same as (B) for male samples.

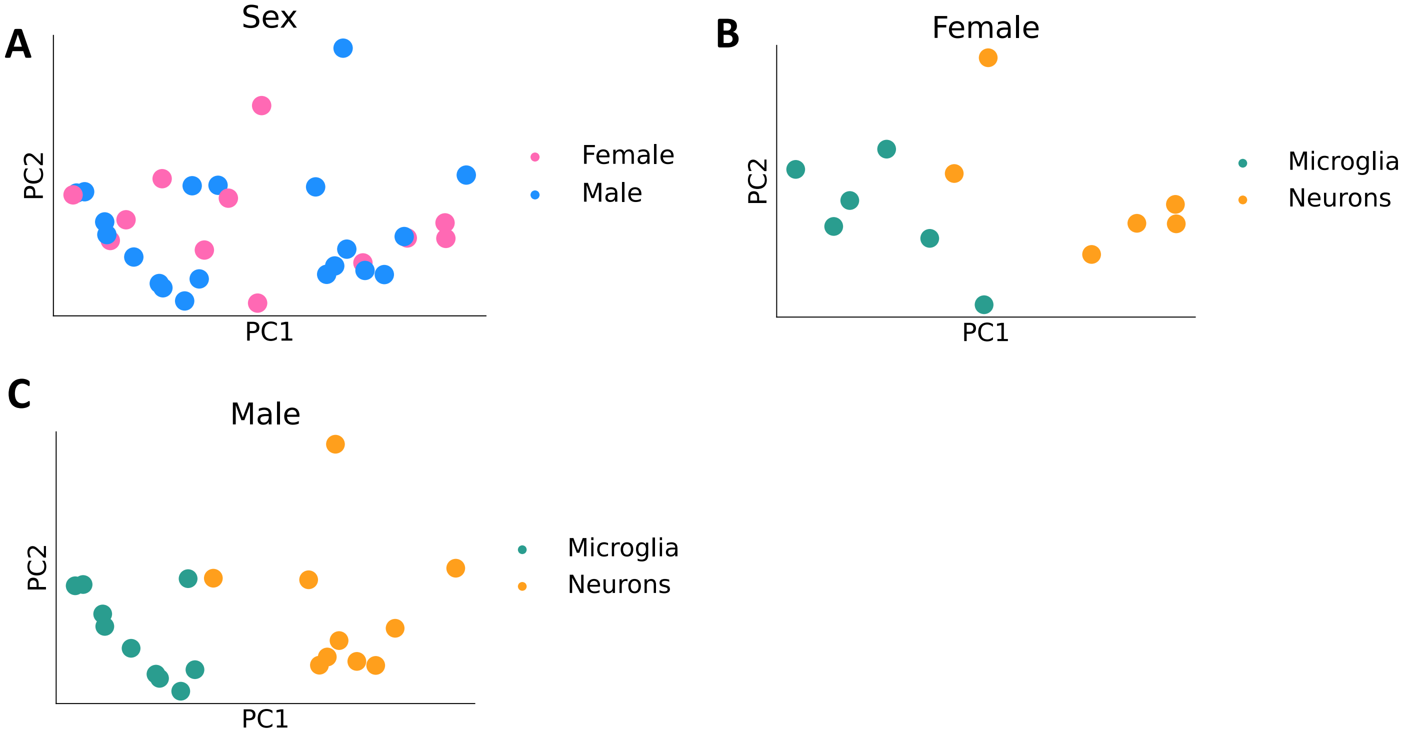

**Supplementary Figure 6: Differential expression of tRFs between neurons and microglia from live human brain**

A) Volcano plot showing tRFs with altered levels in neurons and microglia (in orange – tRFs enriched in neurons, in green – tRFs enriched in microglia). C) Neuron and microglia specific levels of tRFs originating from the tRNAs of origin of tRFs DE in Klotho KO. On the left – tRNA of origin of all three tRFs upregulated in Klotho KO (trna26_AsnGTT_1), on the right – three tRNAs of origin of tRFs downregulated in Klotho KO (tRNA1_LeuAAG_14, tRNA27_LeuTAG_16, tRNA42_LeuTAG_17); **: $p_{adj}$<=0.01, ns – non-significant.

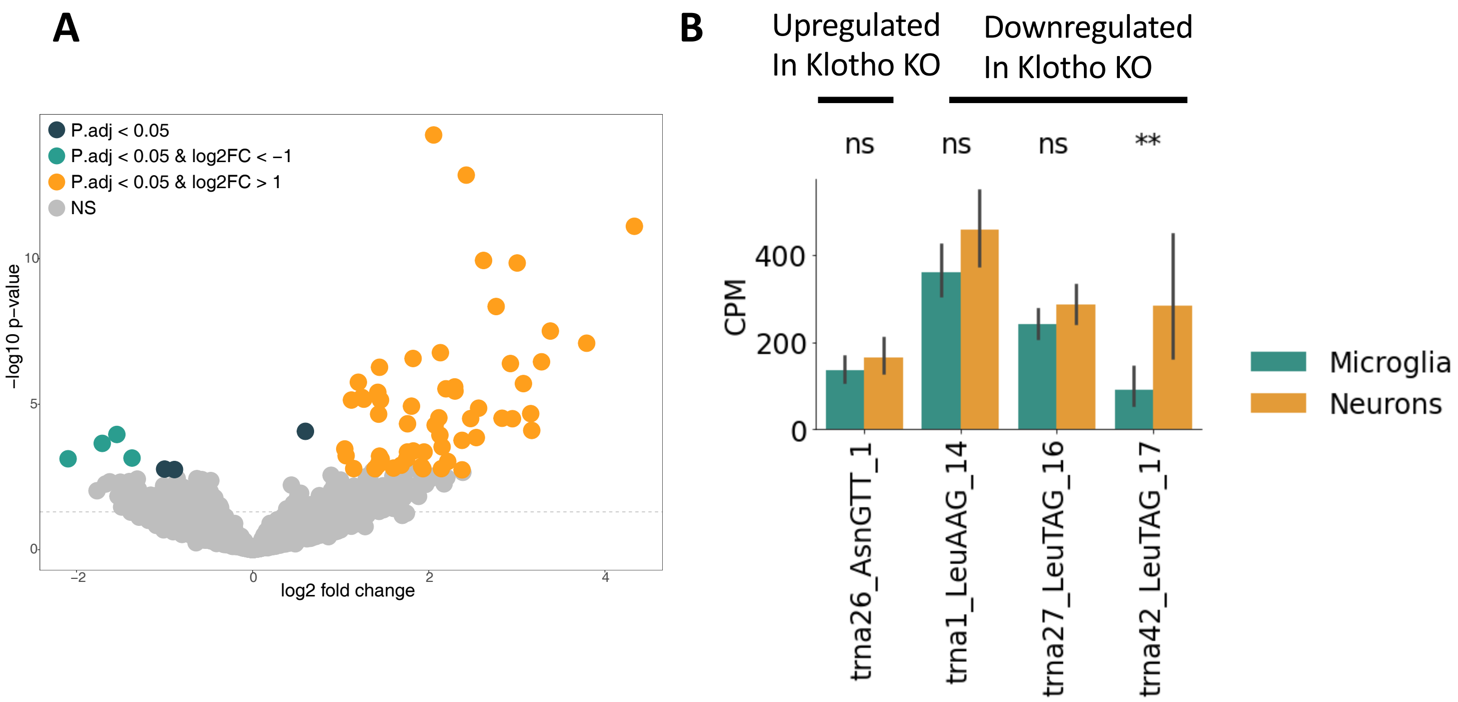
